# Alcohol dependence activates ventral tegmental area projections to central amygdala in male mice and rats

**DOI:** 10.1101/2020.11.02.365445

**Authors:** Elizabeth M Avegno, Chelsea R Kasten, William B Snyder, Leslie K Kelley, Thomas D Lobell, Taylor J Templeton, Michael Constans, Tiffany A Wills, Jason W Middleton, Nicholas W Gilpin

## Abstract

The neural adaptations that occur during the transition to alcohol dependence are not entirely understood, but may include a gradual recruitment of brain stress circuitry by mesolimbic reward circuitry that is activated during early stages of alcohol use. Here, we focused on dopaminergic and non-dopaminergic projections from the ventral tegmental area (VTA), important for mediating acute alcohol reinforcement, to the central nucleus of the amygdala (CeA), important for alcohol dependence-related negative affect and escalated alcohol drinking. The VTA projects directly to the CeA, but the functional relevance of this circuit is not fully established. Therefore, we combined retrograde and anterograde tracing, anatomical, and electrophysiological experiments in mice and rats to demonstrate that the CeA receives input from both dopaminergic and non-dopaminergic projection neurons primarily from the lateral VTA. We then used slice electrophysiology and fos immunohistochemistry to test the effects of alcohol dependence on activity and activation profiles of CeA-projecting neurons in the VTA. Our data indicate that alcohol dependence activates midbrain projections to the central amygdala, suggesting that VTA projections may trigger plasticity in the CeA during the transition to alcohol dependence and that this circuit may be involved in mediating behavioral dysregulation associated with alcohol dependence.

## Introduction

Alcohol use disorder (AUD) affects over 14 million Americans^1^, costing the US ~$249 billion annually^2^. The neural adaptations that occur during the transition to alcohol dependence include a switch from primary activation of mesolimbic circuitry, important for mediating positive reinforcing effects of alcohol, to a gradual recruitment of stress circuitry, important for mediating withdrawal-associated negative affect and negative reinforcement^3^. The ventral tegmental area (VTA) is a midbrain source of dopamine (DA) neurons, although only approximately 55-65% of VTA neurons are dopaminergic, with an estimated 30-35% GABAergic and 2-5% glutamatergic neurons^4–7^. VTA DA neuron function mediates positive reinforcing effects of alcohol in non-dependent rats^8^. Chronic high-dose alcohol exposure produces dependence in rats^9^ and mice^10^; this procedure also alters central amygdala (CeA) synaptic transmission^11^ and neuropeptide signaling^12^ that drive negative affect and escalated alcohol drinking during alcohol withdrawal^13^. The VTA projects directly to the CeA^14–16^, but the functional relevance of this circuit is not fully understood, and no study has examined the effect of chronic alcohol on this circuit.

The VTA mediates the acute positive reinforcing effects of alcohol^17^, and chronic alcohol and drug exposure each produce VTA neuroadaptations^18–21^. VTA neurons are often subdivided based on their projection target and molecular signature (e.g., those that release DA, glutamate, or GABA solely, or co-release two neurotransmitters), and these neuronal subpopulations exhibit differential responses to rewarding and aversive stimuli^22,23^. VTA neurons exhibit varied responses to acute and sub-chronic alcohol, with a subset of medially-located DA neurons demonstrating increased excitatory responses to alcohol relative to lateral VTA DA or non-DA neurons^19,24^. It is not known how chronic alcohol exposure produces circuit-specific plasticity of VTA neurons in alcohol-dependent animals. Here, we demonstrate alcohol withdrawal-induced activation of CeA-projecting VTA neurons in alcohol-dependent rats and mice.

In contrast to the VTA, which is often studied in the context of acute response to EtOH (e.g.,^25^), studies on CeA synaptic transmission and cellular activation have focused more on chronic EtOH exposure (although non-dependent binge-like drinking has also been shown to engage this brain region; e.g.^26,27^). Chronic alcohol exposure increases GABAergic transmission in the CeA of alcohol-dependent rats during withdrawal^11^. Chronic alcohol exposure also increases glutamatergic transmission in dependent rats^28^ and expression of proteins involved in glutamatergic signaling pathways in the CeA of non-dependent mice^29^. These alcohol effects on CeA synaptic transmission and neuropeptide signaling likely mediate withdrawal-induced increases in anxiety-like behavior and alcohol drinking in alcohol-dependent rats^13^. One prior study implicated VTA-CeA projections in anxiety-like behavior by showing that transgenic mice with hyperactive VTA neurons exhibit high anxiety-like behavior, a phenomenon that is reversed by site-specific administration of a D1 receptor antagonist in the CeA, presumably via blockade of midbrain inputs to the CeA^30^.

The VTA sends direct projections to CeA^14–16^, but it is not known how alcohol affects cellular activity in this circuit. Much of what has been inferred about the function of this circuit has come from site-specific modulation of local DA within the CeA, with the assumption that DA in the CeA is released from VTA terminals (e.g.,^30,31^). Here, we sought to more thoroughly characterize the location, function and expression profiles of CeA-projecting VTA neurons in mice and rats. We describe a population of CeA-projecting midbrain neurons and demonstrate activation of the VTA-CeA circuit in both alcohol-dependent mice and rats during withdrawal. Our results indicate that alcohol dependence activates midbrain projections to the amygdala, suggesting that VTA projections may trigger plasticity in the CeA during the transition to alcohol dependence and that this circuit may be involved in mediating alcohol dependence-associated behavioral dysregulation.

## Materials and Methods

### Animals

All procedures were approved by the Institutional Animal Care and Use Committee of the Louisiana State University Health Sciences Center, and were carried out in accordance with the National Institute of Health guidelines. Adult male Long Evans rats (Charles River, Raleigh, NC, USA) weighing ~300 g at start of experiments were pair-housed in a humidity- and temperature-controlled (22°C) vivarium on a 12-hr light/dark cycle (lights off at 8:00 am), with *ad libitum* access to food and water. Adult male C57Bl/6J mice (Jackson Laboratories, Bar Harbor, ME, USA) were received at 10 weeks of age. Mice were housed 2-3/cage with *ad libitum* access to food and water under a 12:12 hr light-dark cycle (lights off at 6:00 pm). All animals were acclimated to housing conditions and light-dark cycles for 1 week before start of experiments. A total of 34 rats and 16 mice were used across experiments.

### Stereotaxic Surgeries

Rats were anesthetized with isoflurane and mounted into a stereotaxic frame (Kopf Instruments, Tujunga, CA, USA). For retrograde tracing experiments, 0.5 μL Retrobeads (Lumafluor, Naples, FL, USA) or Alexa Fluor-488 labeled cholera toxin B (CTxB; ThermoFisher; Waltham, MA, USA) were injected bilaterally into the CeA, using the following coordinates from Bregma: −2.5 mm posterior, ±4.0 mm lateral, −8.4 mm ventral. For viral experiments, rats received 0.5 μL bilateral injections of either channelrhodopsin virus (AAV5-hSyn-hChR2-mCherry, lot AV4320C, UNC Vector Core, Chapel Hill, NC, USA), or control virus (pAAV-hSyn-EGFP, lot v5484) into the VTA, using the following coordinates from Bregma: −5.75 mm posterior, ±0.75 mm lateral, −8.0 mm ventral. Viral titers in all experiments were between 4.1 and 4.6 x 10^12^ viral molecules/mL virus. After all surgical procedures, rats were monitored to ensure full recovery from anesthesia before being singly housed for one night. Rats were treated with the analgesic flunixin (2.5 mg/kg, subcutaneous) once immediately prior to surgery and once every 12 hours for 24 hours post-op. Rats were treated with the antibiotic cefazolin (20 mg/kg, intramuscular) once just prior to surgery and once 24 hours post-op to prevent infection. Weight and appearance were also monitored for a period of 7 days following surgery to detect any signs of distress or post-operative complications.

Surgical procedures in mice were modified from those of Kasten & Boehm (2014)^32^. Mice were anesthetized using a ketamine/xylazine cocktail (1 mL ketamine, 0.1 mL xylazine, and 8.9 mL sterile saline, i.p.) and mounted into a stereotaxic frame (Leica Biosystems, Buffalo Grove, IL, USA). Mice received 0.25 μL bilateral injections of either channelrhodopsin or control virus into the VTA using the following coordinates from Bregma: −3.64 mm posterior, ±0.5 mm lateral, −4.2 mm ventral. Mice were treated with the analgesic ketoprofen (5 mg/mL, subcutaneous) once during surgery, then again approximately 24 hrs and 48 hrs post-surgery. Weight and appearance were monitored for 7 days following surgery to detect any signs of distress or post-operative complications. All drugs were given at a volume of 0.1 mL per 10 g of body weight.

### Chronic Intermittent Exposure to Alcohol Vapor

Chronic intermittent exposure (CIE) to alcohol vapor was used to induce dependence in rats and mice^33^. Rats were pair-housed in sealed chambers (La Jolla Alcohol Research Inc., La Jolla, CA, USA) and exposed to alcohol vapor for 14 hr/day (Fig. 3A). Blood alcohol levels (BALs) were measured 1-2 times weekly and analyzed using an Analox AM1 analyzer (Analox Instruments, LTD, Lunenberg, MA, USA). Rats were maintained within a BAL range of 150-250 mg/dL over the course of a minimum of 4 weeks (mean 180 ± 17.0 mg/dL across experiments). All stereotaxic injections were given prior to initiation of CIE to minimize potential interactions between alcohol and the anesthetic isoflurane.

Mice were group housed and exposed to alcohol for 16 hr/day for four consecutive days. Prior to vapor exposure, mice were given a daily injection of either pyrazole (air control, 1 mmol/kg) or pyrazole + ethanol (ethanol group, 1 mmol/kg + 0.8 g/kg, respectively) to impair the metabolism of ethanol. Trunk blood was obtained to spot check BALs (mean 202 ± 21.4 mg/dL across experiments). Mice received two full cycles of vapor exposure (4 days on, 3 days off), and were sacrificed for electrophysiological recordings either one hour (“intoxication”) or five hours (“withdrawal”) following the end of the fourth day of vapor exposure during their second cycle of exposure (Fig. 4A). Trunk blood was collected during sacrifice; mean BACs taken from mice were 190 ± 23 mg/dL (“intoxication”) and 38 ± 16 mg/dL (“withdrawal”). Mice were injected with virus and given approximately 6.5 weeks to allow the virus to travel from the VTA to CeA terminals prior to the start of CIE.

### In vitro Electrophysiology

CIE rats (N = 5) were sacrificed 6-8 hr after cessation of vapor exposure, when BALs were at or near zero (mean 22.4 ± 13.4 mg/dL). Under isoflurane anesthesia, rats were transcardially perfused with room temperature (~25°C) NMDG artificial cerebrospinal fluid (aCSF) containing the following (in mM): 92 NMDG, 2.5 KCl, 1.25 NaH_2_PO_4_, 30 NaHCO_3_, 20 HEPES, 25 glucose, 2 thiourea, 0.5 CaCl_2_, 10 MgSO_4_·7 H_2_O, 5 Na-ascorbate, 3 Na-pyruvate. 300 μm-thick coronal sections containing the CeA or VTA were collected using a vibratome (Leica VT1200S, Nussloch, Germany). Sections were incubated in NMDG aCSF at 37°C for 12 min, then transferred to a room temperature holding aCSF solution containing the following (in mM): 92 NaCl, 2.5 KCl, 1.25 NaH_2_PO_4_, 30 NaHCO_3_, 20 HEPES, 25 glucose, 2 thiourea, 2 CaCl_2_, 2 MgSO_4_·7 H_2_O, 5 Na-ascorbate, 3 Na-pyruvate. Mice were sacrificed under isoflurane anesthesia, and 300 μm-thick coronal sections containing the CeA were collected as described above. Slices were allowed to recover for one hour prior to recording. Slices were visualized with oblique infrared light illumination, a w60 water immersion objective (LUMPLFLN60X/W, Olympus, Tokyo, Japan) and a CCD camera (Retiga 2000R, QImaging, Surrey, BC, Canada). Data were sampled at 10 kHz and Bessel filtered at 4 kHz using an acquisition control software package Ephus^34^.

Recording conditions were identical for mice and rats. Sections were transferred to a recording aCSF solution containing the following (in mM): 130 NaCl, 3.5 KCl, 2 CaCl_2_, 1.25 NaH_2_PO_4_, 1.5 MgSO_4_·7 H_2_O, 24 NaHCO_3_, 10 glucose. Recording aCSF was maintained at 32-34°C using an in-line heater (Warner Instruments, Hamden, CT). For VTA recordings, retrobead-containing (CeA-projecting) neurons were identified by their fluorescence using a filter set (Chroma, Bellows Falls, VT, USA). Recordings were performed using an internal recording solution containing the following (in mM): 140 K-gluconate, 5 KCl, 0.2 EGTA, 10 HEPES, 2 MgCl_2_·6 H_2_O, 4 Mg-ATP, 0.3 Na_2_-GTP, 10 Na_2_-phosphocreatine (pH 7.2-7.3, 285-295 mOsm). For CeA optical stimulation experiments, voltage clamp recordings were performed with cells clamped at −75 mV. Light pulses (~13 mW/mm^2^ power density, 0.5 ms pulse duration, 5 Hz pulse frequency) were delivered, and an average of ten traces were recorded for each cell’s baseline response. For those cells in which an optically-evoked postsynaptic current (ePSC) was observed, SCH 23390 hydrochloride (5 μM; R&D Systems, Minneapolis, MN, USA) was added to the aCSF, and an additional 5 traces were recorded following 5 minutes of incubation to determine the effect of D1 receptor blockade on ePSCs. 2,3-Dioxo-6-nitro-1,2,3,4-tetrahydrobenzo[f]quinoxaline-7-sulfonamide (NBQX; 10 μM; R&D Systems) was then added to determine the effect of AMPA receptor blockade on ePSCs. Liquid junction potentials were not corrected during recordings. Experiments with a series resistance >30 MΩ or a >20% change in series resistance were excluded from analysis.

### TH and Fos Immunohistochemistry in Rats

Under isoflurane anesthesia, rats (n = 5 CIE and 5 naïve) were transcardially perfused with saline followed by 4% paraformaldehyde (PFA). After perfusion, brains were post-fixed for 24 hr, transferred to 30% sucrose solution for 24-48 hr, then snap-frozen in isopentane. 30 μm-thick coronal sections were collected and stored in a 0.1% sodium azide in 1X phosphate buffered saline (PBS) until processing.

Sections were incubated in 3% H_2_O_2_ for 5 min, followed by two 10-min washes PBS. Slices were incubated in a blocking solution containing 1% BSA, 10% normal donkey serum, and 0.3% Triton-X 100 in PBS for 1 hr before a 48-hr incubation at 4°C in 1:1,000 mouse anti-TH (Millipore; St. Louis, MO, USA) and 1:3,000 rabbit anti-cFos (Millipore), with 1% BSA, 2% normal donkey serum, and 0.3% Triton-X 100. Following primary antibody incubation, slices were transferred to a solution with 1:500 AlexaFluor 647-conjugated donkey anti-mouse (ThermoFisher) and 1% normal donkey serum in TNB buffer (Perkin Elmer; Waltham, MA, USA) for 30 min, then transferred to a solution with 1:500 AlexaFluor 647-conjugated donkey anti-mouse (ThermoFisher) and 1% normal donkey serum in ImmPRESS horse anti-rabbit solution (Vector Laboratories; Burlingame, CA, USA) for 30 min. Sections were transferred to four 5-min washes in TNT buffer, then incubated in TSA plus cyanine 3 (Perkin Elmer) for 10 min. Following two 10-min washes in TNT buffer, sections were mounted and coverslipped with Fluoromount (Sigma).

To quantify Retrobead, fos, and TH expression, three midbrain sections were analyzed per animal at the same approximate bregma depths (anterior, mid, and posterior). 20x images of the entire midbrain were collected, then stitched together to reconstruct the full midbrain. Retrobead+ neurons were manually counted, then assessed for overlapping co-expression of fos and/or TH. Boundaries of the VTA and SN on the stitched midbrain image were drawn using the Paxinos and Watson atlas (2007)^35^ as a guide, and only neurons within the boundaries were counted. Accurate injection site was verified for each rat, and only those with Retrobead injections inside of the CeA were included for analysis.

### In-Situ Hybridization in Rats

One week following intra-CeA injection of CTxB-488, rats (N = 4) were placed under isoflurane anesthesia and transcardially perfused with saline followed by 4% PFA. After perfusion, brains were post-fixed for 24 hr, transferred to serial concentrations of sucrose (10, 20, and 30%), then snap-frozen in isopentane. 10 μm-thick VTA-containing coronal sections were collected and stored at −20°C in a cryoprotectant solution (30% ethylene glycol, 30% glycerin, 2% DMSO, 30% sucrose in PBS) until processing. Sections were prepared for processing by 10 min washes in PBS and TBS, then quenched in a 3% H_2_O_2_ in TBST solution (20 min), followed by a 3% H_2_O_2_ in TBS solution (40 min). After two 10 min washes in TBS, slices were mounted to a glass slide, allowed to air dry overnight, then processed according to manufacturer instructions (Bio-Techne; Newark, CA, USA).

Slides were dehydrated in 100% ethanol for 2 min, then incubated in Protease III for 30 min. Slides were washed twice with agitation in PBS for 2 min, then incubated for 2 hr at 40 °C in a solution containing either probes for *Th* (which transcribes TH, a marker for dopaminergic neurons; Rn-Th-C3; lot20177A) and *Slc17a6* (which transcribes vesicular glutamate transporter 2 [vGluT2], a marker for glutamatergic neurons; Rn-Slc17a6-C2; lot 181550), or, on a separate set of tissue, probes for *Th* (Rn-Th-C2; lot 201773G) and *Slc32a1* (which transcribes vesicular inhibitory amino acid transporter [VIAAT], a marker for GABAergic neurons; Rn-Slc32a1-C3; lot 20077B). Following a series of amplification steps, slides were coverslipped with ProLong Gold Antifade Reagent with DAPI (ThermoFisher) and imaged within 48 hours.

### VTA Terminal Imaging in Rats and Mice

Under isoflurane anesthesia, rats (N = 3) or mice (N = 3) were transcardially perfused with saline followed by 4% PFA. 30 μm-thick coronal sections were collected as described above. Sections were washed in PBS-T for 10 min, followed by two 5-min washes in PBS. Sections were then incubated in 1:100 NeuroTrace™ 530/615 Red Fluorescent Nissl Stain (ThermoFisher) in PBS for 2 hr and subsequently washed in PBS-T for 10 min, followed by two 5-min washes in PBS. Following an overnight incubation in PBS at 4°C, sections were mounted and coverslipped with ProLong Gold antifade mountant with DAPI (ThermoFisher).

### Imaging

Images of Retrobead containing midbrain neurons and pAAV-hSyn-EGFP-containing terminals in CeA were obtained with a fluorescent/brightfield microscope (Olympus, Tokyo, Japan). Images of CeA-projecting VTA neurons from the *in-situ* hybridization experiment were obtained with a Keyence BZ-X810 all-in-one fluorescent microscope (Keyence Corporation; Osaka, Japan).

### Experimental Design and Statistical Analysis

Statistical analysis was performed in Prism 7.03 (GraphPad Software, La Jolla, CA, USA). Data were analyzed using two-tailed *t*-test, one-way ANOVA, and Fisher’s exact test. A *p* value of < 0.05 was considered significant. All tests and variables are identified in the Results section.

## Results

### Heterogeneous population of midbrain neurons project to CeA

To characterize CeA-projecting VTA neurons in the naïve brain, we stereotaxically injected Retrobeads into the CeA of adult male rats (Fig. 1A) and performed immunohistochemistry on VTA-containing tissue sections (Fig. 1B-C). We found that CeA-projecting VTA neurons are primarily located within the lateral VTA, with fewer CeA-projecting medial VTA neurons (Fig. 1D). Only 35.5 (± 7.1)% of neurons expressed TH, a marker for dopaminergic neurons, indicating a substantial population of putative glutamatergic or GABAergic neurons. We also observed a population of CeA-projecting neurons located in the substantia nigra, of which 24.8 (± 4.6)% co-expressed TH. CeA-projecting midbrain neurons were located throughout the anterior-posterior axis.

**Figure 1:**
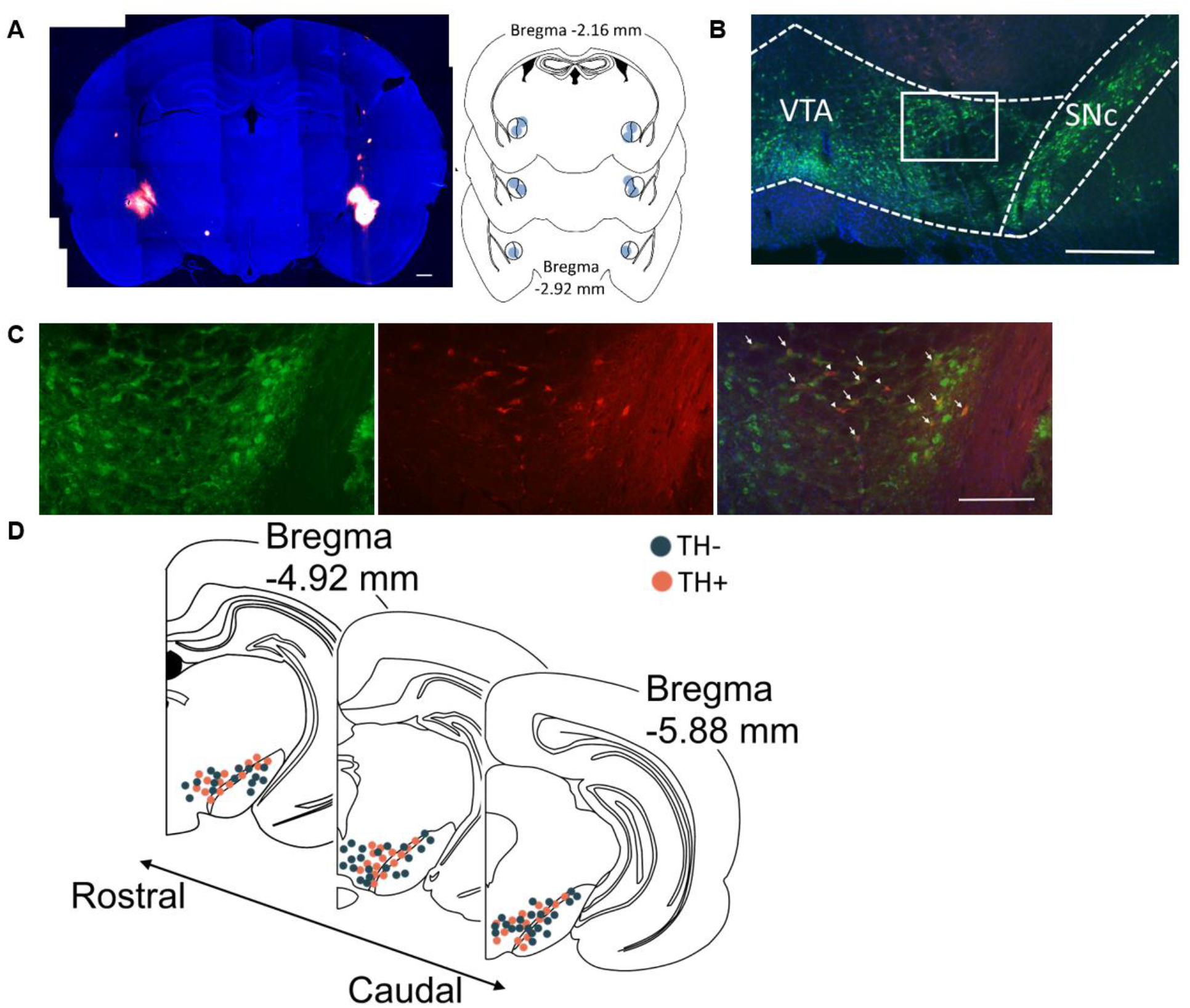
Protein analysis demonstrates that a mixed population of VTA neurons project to CeA of naïve rats. **A**, 4x tiled image of retrograde tracer (red) injected into CeA of adult male rat. Nuclei are stained with DAPI and appear blue. Scale bar, 200 μm. Inset shows center of CeA injection sites for rats used in experiment. **B**, 4x and **C**, 20x images of CeA-projecting (red) neurons in midbrain. Co-expression of TH, as detected by immunohistochemistry, shown in green. Arrows denote CeA-projecting TH+ VTA neurons; arrowheads denote CeA-projecting TH-neurons. Scale bar, 200 μm (**B**), 50 μm (**C**). **D**, Representative location of CeA-projecting neurons within the midbrain of rat, across the anterior-posterior axis. TH+ neurons indicated in red-orange; TH-neurons indicated in dark blue. Data averaged from 5 rats (n = 3 sections/rat).

Because we observed a substantial population of TH-negative CeA-projecting VTA neurons, we next performed *in-situ* hybridization studies to further evaluate the expression profile of these cells. Alexa Fluor-488 labeled cholera toxin B (CTxB) was stereotaxically injected into the CeA of adult male rats (Fig. 2A), and RNAscope was performed on VTA-containing sections to evaluate expression of *Th* (which transcribes TH) and *Slc17a6* (which transcribes vesicular glutamate transporter 2 [vGluT2], a marker for glutamatergic neurons; Fig. 2B–2C). In a separate set of sections from the same animals, expression of *Th* and *Slc32a1* (which transcribes vesicular inhibitory amino acid transporter [VIAAT], a marker for GABAergic neurons) was evaluated (Fig. 2B & 2D). In the first set of tissue, we observed a similar percentage of *Th*-positive CeA-projecting VTA neurons (27.8 ± 3.5%) relative to that observed from our immunohistochemical analysis. Of those *Th*-positive CeA-projecting VTA neurons, 51% co-expressed the glutamatergic marker *Slc17a6*. A separate 27.3 (± 5.8)% of neurons expressed solely *Slc17a6*, indicating a large proportion of CeA-projecting VTA neurons capable of glutamate release, about half of which are dopaminergic. In a separate set of tissue, 31.5 (± 6.4)% of CeA-projecting VTA neurons expressed the GABAergic marker *Slc32a1*, with only 0.5% of neurons co-expressing *Th* and *Slc32a1*, suggesting that a substantial number of CeA-projecting VTA cells are GABAergic, but that very few CeA-projecting VTA neurons co-express dopamine and GABA. These data, summarized in Fig. 2B & 2E-F, indicate that CeA-projecting VTA neurons have diverse molecular phenotypes that are intermixed along the medial-lateral and anterior-posterior axes.

**Figure 2:**
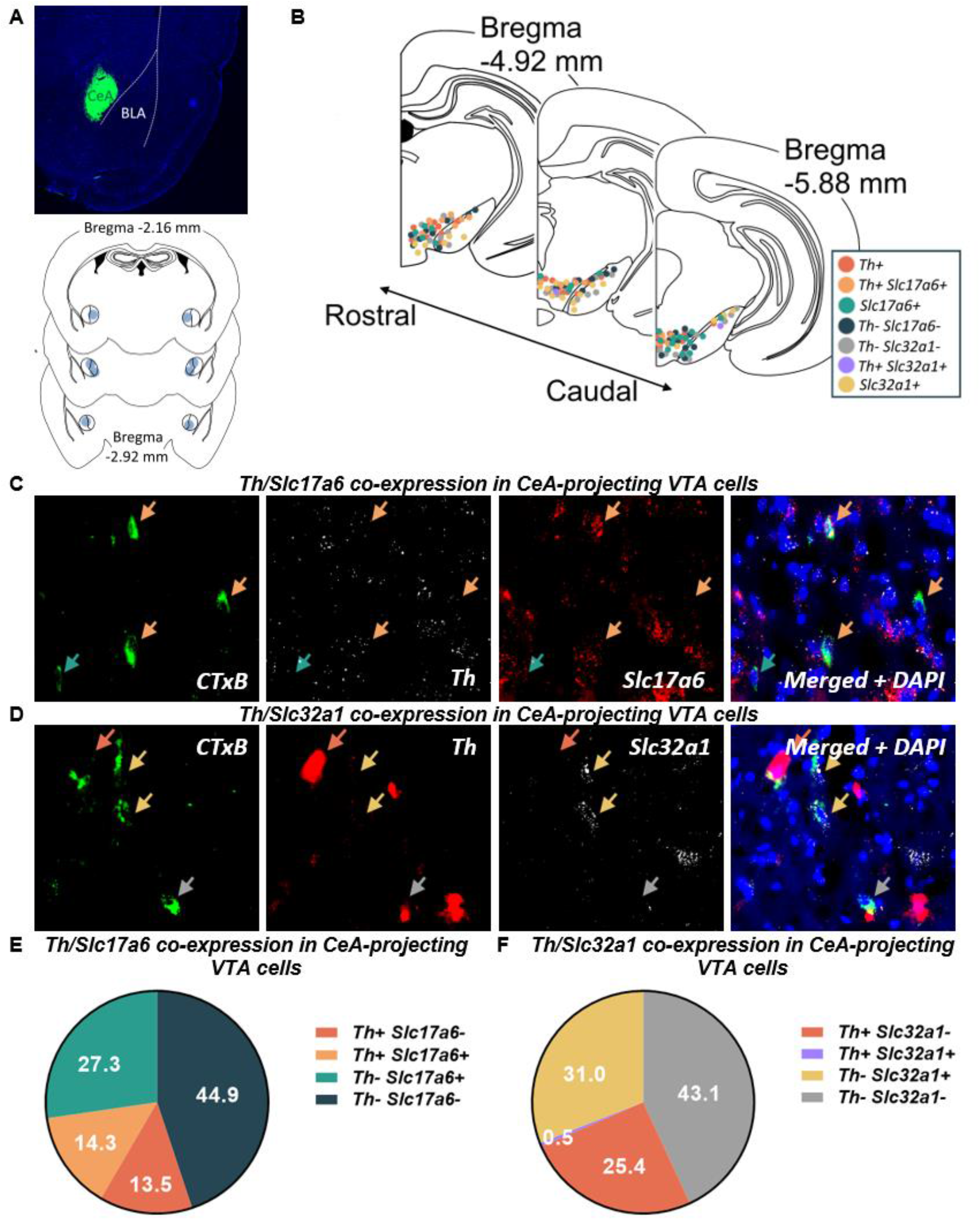
RNA expression analysis demonstrates that a mixed population of VTA neurons project to CeA of naïve rats. **A**, 4x tiled image of retrograde tracer (CTxB, green) injected into CeA of adult male rat. Nuclei are stained with DAPI and appear blue. CeA, central amygdala; BLA, basolateral amygdala. Inset shows center of CeA injection sites for rats used in experiment. **B**, Representative expression profile of CeA-projecting neurons within the midbrain of rat, as determined by *in situ* hybridization experiments, across the anterior-posterior axis. Expression of only one marker (*Th*, red-orange; *Slc17a6*, green; *Slc32a1*, yellow), co-expression of two markers *(Th+ Slc17a6+*, orange; *Th*+ *Slc32a1*+, purple), or absence of markers (*Th-Slc17a6*-[putative GABAergic], dark blue; *Th*-*Slc32a1*-[putative glutamatergic], grey) indicated. Data averaged from 4 rats (n = 3 sections/rat). **C**, 40x images of CeA-projecting (green) neurons in midbrain. Co-expression of *Th* (white) and *Slc17a6* (red) shown. Orange arrows denote CeA-projecting *Th*+ *Slc17a6*+ VTA neurons; green arrow denotes CeA-projecting *Th-Slc17a6*+ neuron. **D**, 40x images of CeA-projecting (green) neurons in midbrain. Co-expression of *Th* (red) and *Slc32a1* (white) shown. Red-orange arrow denotes CeA-projecting *Th*+ *Slc32a1*-VTA neuron; yellow arrows denote CeA-projecting *Th-Slc32a1*+ VTA neurons; grey arrow denotes CeA-projecting *Th-Slc32a1*-neuron. Expression profiles from neurons represented in **C** and **D** quantified in **E** and **F**, respectively.

To map VTA inputs into the CeA, we combined viral injections in the VTA with imaging of terminals in the CeA. We injected pAAV-hSyn-EGFP bilaterally into the VTA of mice and rats to transfect all neurons with GFP (Fig. 3A), and GFP-containing terminals were imaged in the CeA (Fig. 3B). We observed VTA inputs into both the medial and lateral subdivisions of the CeA in the rat, across the anterior-posterior axis (Fig. 3C). Interestingly, we observed slight species-specific differences in the circuit; in the mouse, VTA input was restricted primarily to the medial and anterior subdivisions of the CeA (Fig. 3D). To assess a functional connection between the two regions, we next injected AAV5/hSyn-Channelrhodopsin-mCherry (ChR2) into the VTA of naïve mice and rats and performed whole cell recordings of CeA neurons. We sampled neurons located near mCherry-containing terminals, and recordings were primarily performed in the medial subdivision of the CeA. In the presence of blue light stimulation, we observed optically-evoked postsynaptic currents (ePSCs) in a subset of neurons (Fig. 3E). Bath application of postsynaptic receptor blockers had a significant effect on ePSC amplitude (F(2,19) = 3.71, *p* = 0.0435; one-way ANOVA). Post-hoc analysis revealed application of D1 receptor antagonist SCH 23390 did not affect the amplitude of ePSCs (78.7 ± 9.5% of baseline; *p* = 0.94, Tukey’s multiple comparisons test), suggesting that activity of D1 receptors does not modulate this postsynaptic response. No association between CeA location and response to SCH 23390 was observed. Addition of the AMPA receptor antagonist NBQX abolished optically-evoked postsynaptic responses in all neurons tested (amplitude 11.3 ± 10.4% of baseline; *p* = 0.048, Tukey’s multiple comparisons test), indicating release of glutamate from VTA terminals into the CeA. Interestingly, we observed ePSCs more readily in the mouse (4 of 12 neurons, 33%) than in the rat (1 of 13 neurons, 8%), although this difference was not statistically significant (*p* = 0.16, Fisher’s exact test; Fig. 3F-G). No significant difference was observed in the resting membrane potential of CeA neurons from mice (−56.5 ± 3.0 mV) and rats (−52.8 ± 2.4 mV; *t*(23) = 0.98; *p* = 0.34; two-tailed *t*-test). Similarly, no difference was observed in baseline firing rate of CeA neurons from mice (1.3 ±1.0 Hz) and rats (3.2 ± 1.3 Hz; *t*(10) = 0.93; *p* = 0.38; two-tailed *t*-test), nor in the proportion of spontaneously active CeA neurons from mice (4 of 12 neurons, 33.3%) and rats (8 of 13 neurons, 61.5%; *p* = 0.24, Fisher’s exact test) was observed.

**Figure 3:**
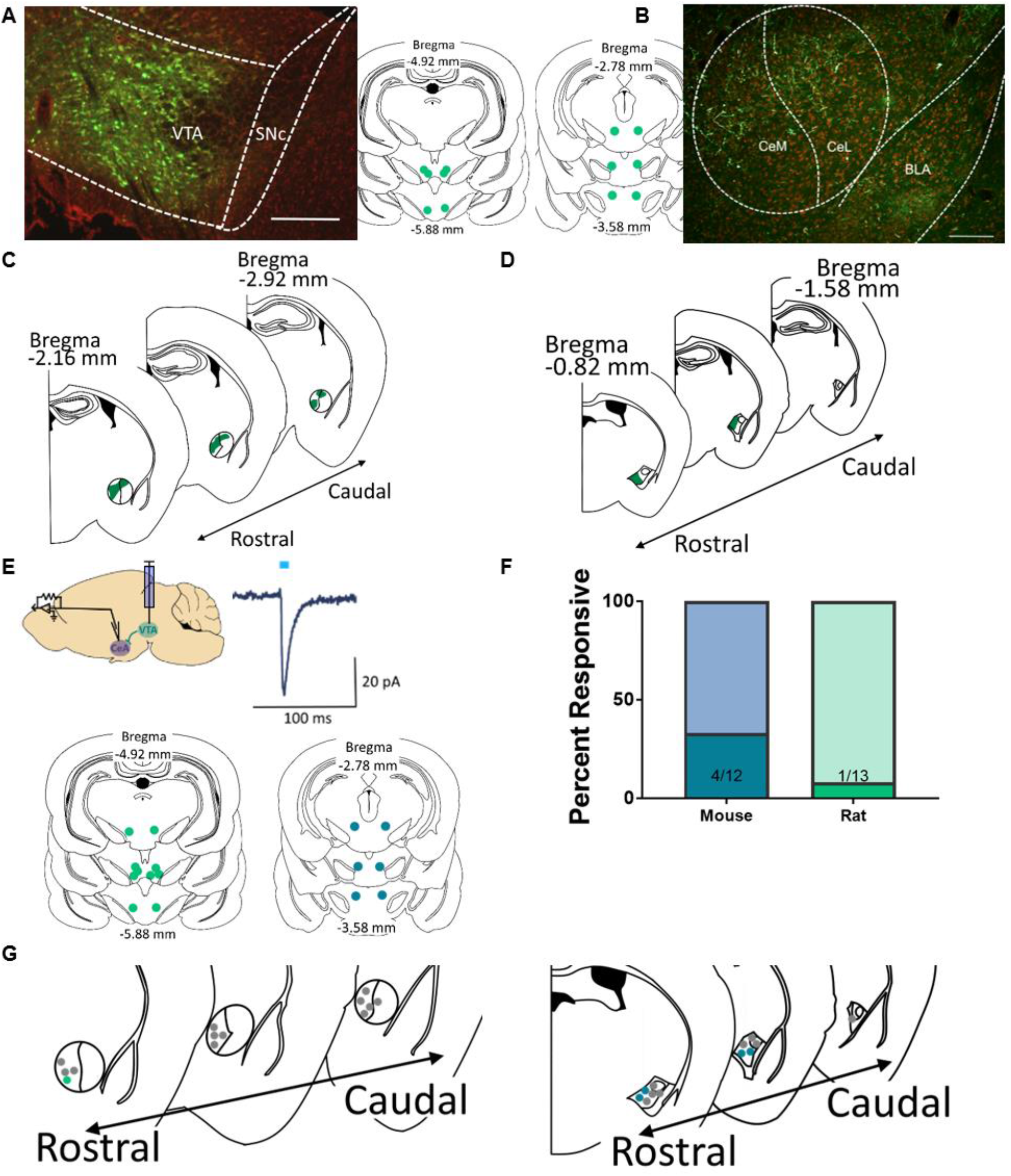
VTA neurons synapse onto CeA neurons in mice and rats. **A**, Representative 4x image of GFP (pAAV-hSyn-EGFP)-expressing neurons in VTA of rat. Neurons are counterstained with red Nissl. Scale bar, 200 μm. Inset shows center of VTA injection sites for rats (left) and mice (right) used in experiment. **B**, 20x image of GFP-containing terminals in the CeA. CeM, medial subdivision of CeA; CeL, lateral subdivision of CeA; BLA, basolateral amygdala. Scale bar, 50 μm. **C**, Map indicating approximate location of VTA terminals in the rat CeA, across the anterior-posterior axis. **D**, Map indicating approximate location of VTA terminals in the rat CeA, across the anterior-posterior axis. Data from **C** and **D** averaged from 3 animals each (n = 3 slices/animal). **E**, Representative baseline optically-evoked postsynaptic current (ePSC) from stimulation of ChR2 (AAV5/hSyn-Channelrhodopsin-mCherry)-containing VTA terminals in CeA (inset). Center of VTA injection sites for rats (left, green) and mice (right, blue) used in experiment shown. **F**, Proportion of CeA neurons that respond to optical stimulation in the mouse and the rat. **G**, Location of recorded neurons in rat (left) and mouse (right). Non-responsive neurons are shown in grey; responsive neurons are shown in green (mouse) and blue (rat).

### Alcohol withdrawal activates VTA-CeA circuit in alcohol-dependent mice and rats

We next tested the effect of CIE on VTA-CeA circuit activation in alcohol-dependent mice and rats. To identify CeA-projecting neurons in the VTA, we stereotaxically injected Retrobeads into the CeA of rats prior to the start of CIE (Fig. 4A). Following a minimum of 4 weeks CIE, rats were sacrificed during withdrawal, and brains were used for immunohistochemical or slice electrophysiology experiments.

**Figure 4:**
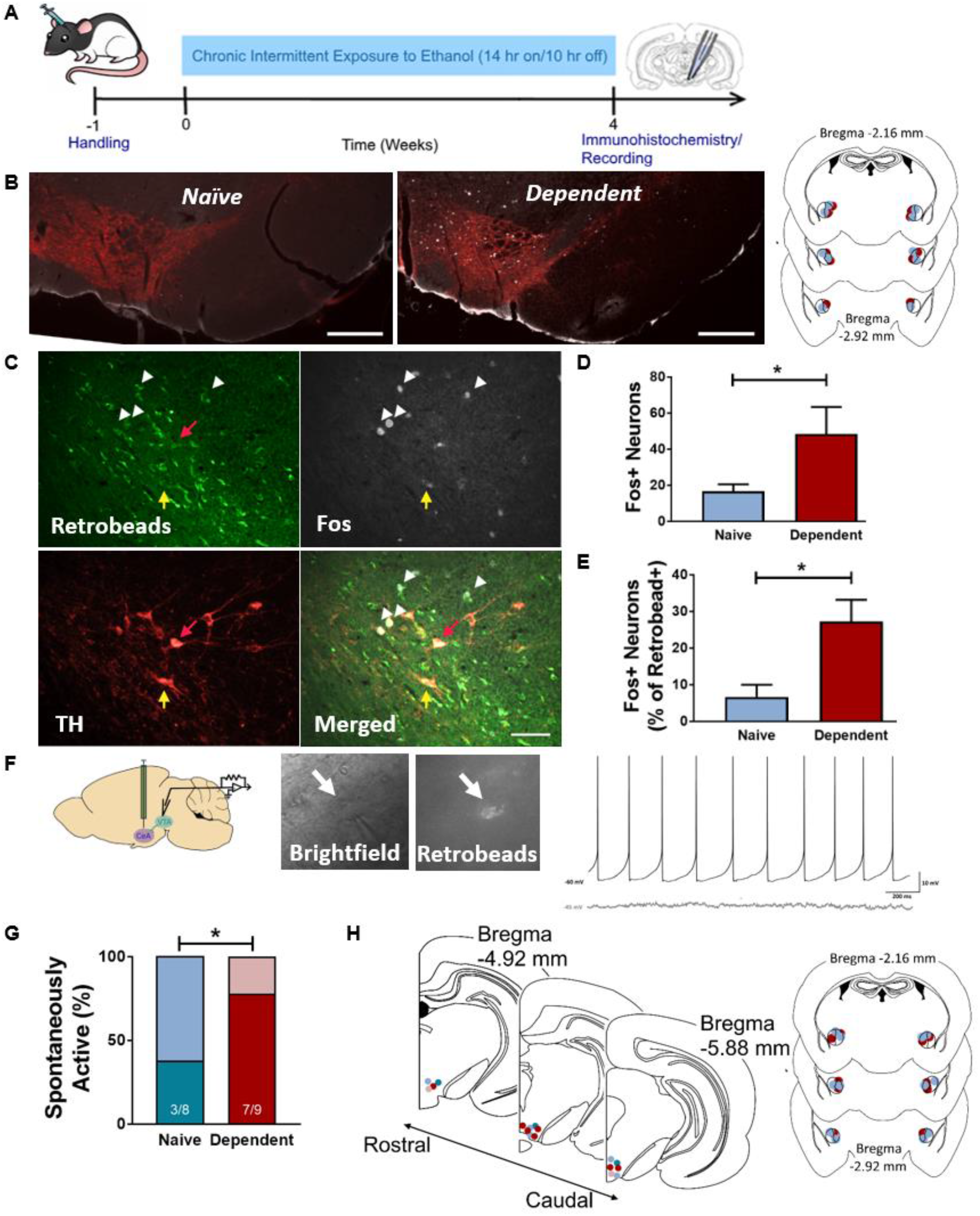
CeA-projecting VTA neurons are activated in alcohol-dependent rats during withdrawal. **A**, Timeline of CIE experiments. **B**, 4x image of midbrain of naïve (left) and alcohol dependent (right) rat. TH shown in red, Fos in white, Retrobeads shown in green. Scale bar, 500 μm. Inset shows center of CeA injection sites for vapor (red) and naïve (blue) rats. **C**, 20x image of VTA of alcohol dependent rat. TH shown in red, Fos in white, Retrobeads shown in green. Scale bar, 100 μm. White arrowheads denote Fos+, retrobead+ neurons; red arrow, TH+, retrobead+ neuron; yellow arrow, Fos+, TH+, retrobead+ neuron. **D**, Total number of Fos+ neurons is significantly greater in VTA of dependent rats (red) compared to naïve controls (blue). **E**, Number of Fos+ neurons as a percentage of all Retrobead+ (CeA-projecting) neurons is significantly greater in VTA of dependent rats (red) compared to naïve controls (blue). (\**p* < 0.05, two-tailed *t*-test. n = 3 sections/rat, 5 rats/group.) **F**, Left, 60x image of Retrobead-containing (CeA-projecting) VTA neuron with patch pipette. Right, representative traces from spontaneously active (black) and silent (grey) CeA-projecting VTA neurons. **G**, Proportion of spontaneously active CeA-projecting VTA neurons of alcohol-dependent rats sacrificed during withdrawal is significantly greater compared to naïve controls. (\**p* < 0.05, Fisher’s exact test.) **F**, Location of recorded neurons in naïve (blue) and alcohol dependent (red) rats. Non-responsive neurons are shown as greyed colors. Inset shows center of CeA injection sites for vapor (red) and naïve (blue) rats. Data in **D**, **E**, and **G** shown as mean ± SEM.

We assessed fos immunoreactivity as an indicator of neuronal activation in the VTA of CIE and control rats (Fig. 4B-C). Alcohol-dependent rats sacrificed during withdrawal exhibited a significantly greater number of total fos+ neurons in the VTA relative to naïve controls (*t*(28) = 2.05, *p* = 0.049, two-tailed *t*-test; Fig. 4D). Alcohol-dependent rats sacrificed during withdrawal also exhibited a significantly greater percentage of fos+ CeA-projecting (Retrobead-containing) VTA neurons relative to naïve controls (*t*(28) = 3.01, *p* = 0.0055, two-tailed *t*-test; Fig. 4E), suggesting potential activation of this population of neurons during withdrawal. Of the Retrobead+ fos+ VTA neurons, 34.0 (± 3.9)% of neurons expressed TH, similar to the overall percentage of CeA-projecting neurons that are TH-positive.

To assess the activity of CeA-projecting VTA neurons from CIE and control rats, whole cell recordings were performed in Retrobead-containing (CeA-projecting) neurons (Fig. 4F). Alcohol-dependent rats sacrificed during withdrawal also exhibited a significantly greater percentage of spontaneously active CeA-projecting VTA neurons (7 of 9 cells, 77.8%) relative to naïve controls (3 of 8 cells, 37.5%; *p* < 0.05, Fisher’s exact test; Fig. 4G-H). No significant difference was observed in the resting membrane potential of CeA-projecting VTA neurons from dependent rats (−49.6 ± 1.4 mV) and naïve controls (−46.9 ± 1.0 mV; *t*(15) = 1.57, *p* = 0.14, two-tailed *t*-test). Within those spontaneously active neurons, we observed no significant difference in baseline firing rate between neurons from dependent rats (2.48 ± 1.8 Hz) and naïve controls (2.16 ± 0.4 Hz; *t*(7) = 0.13, *p* = 0.90, two-tailed *t*-test). We also observed a similar proportion of pacemaking neurons among spontaneously active cells from alcohol-dependent rats (57%) and alcohol-naïve controls (66%). Together with our immunohistochemical data, this suggests that alcohol withdrawal activates CeA-projecting VTA neurons in alcohol-dependent animals.

We next sought to evaluate whether VTA input to the CeA is altered following CIE by combining optogenetics with slice electrophysiology (e.g., Fig. 3E). Here, we used adult male mice because we more readily observed optically evoked PSCs in the mouse than the rat (Fig. 3F). Mice were made dependent on alcohol using CIE (Fig. 5A), then sacrificed for electrophysiological recordings either one hour (“intoxication”, when mean BACs taken from mice were 190 ± 23 mg/dL) or five hours (“withdrawal”, when mean BACs taken from mice were 38 ± 16 mg/dL) after vapor cessation. An intoxication group was included to assess whether cumulative effects of chronic alcohol exposure on VTA-CeA circuit function are evident during intoxication (when BALs are > 0) and/or during withdrawal (when BALs are approaching or equal to zero). We observed optically evoked PSCs in a significantly greater proportion of CeA neurons from alcohol-dependent mice sacrificed during withdrawal (9 of 11 cells, 81.8%) relative to air controls (5 of 18 cells, 27.8%; *p* = 0.0078, Fisher’s exact test; Fig. 5B). Similarly, we observed optically evoked PSCs in a significantly greater proportion of CeA neurons from alcohol-dependent mice sacrificed during intoxication (10 of 13 cells, 76.9%) relative to air controls (5 of 18 cells, 27.8%; *p* = 0.011, Fisher’s exact test; Fig. 5B). We did not observe any significant difference in the mean amplitude of ePSCs from alcohol-dependent mice sacrificed during intoxication (−103.1 ± 29.0 pA), alcohol-dependent mice sacrificed during withdrawal (−83.1 ± 34.0 pA), and alcohol-naïve mice (−37.76 ± 9.8 pA; F(2,21) = 0.98, *p* = 0.39, one-way ANOVA; Fig. 5C). A two-way ANOVA revealed a significant effect of bath-applied drug on ePSC amplitude (F(1.17,21.65) = 20.19, *p* < 0.0001; Fig. 5D). Post-hoc analysis indicates no significant effect of SCH 23390 application (*p* = 0.74 compared to amplitude at baseline), but NBQX significantly decreased ePSC amplitude (*p* = 0.0006 compared to amplitude with SCH 23390 alone; Tukey’s multiple comparisons test). Similar effects of SCH 23390 and NBQX application on ePSC amplitude were observed across groups (F(4,37) = 1.38, *p* = 0.2587, two-way ANOVA). No significant difference was observed in the resting membrane potential of CeA neurons from alcohol-dependent mice sacrificed during intoxication (−58.9 ± 2.7 mV), alcohol-dependent mice sacrificed during withdrawal (−53.2 ± 1.9 mV), and alcohol-naïve mice (−56.1 ± 2.2 mV; F(2,39) = 0.26; *p* = 0.77, one-way ANOVA). A similar proportion of spontaneously active CeA neurons was observed in alcohol-dependent mice sacrificed during intoxication (5 of 11 cells, 45.5%), alcohol-dependent mice sacrificed during withdrawal (9 of 15 cells, 60%), and alcohol-naïve controls (6 of 16, 37.5%). No significant difference was observed in baseline firing rate of CeA neurons from alcohol-dependent mice sacrificed during intoxication (5.7 ± 4.0 Hz), alcohol-dependent mice sacrificed during withdrawal (1.5 ± 0.8 Hz), and alcohol-naïve mice (6.1 ± 3.9 Hz; F(2,14) = 0.68; *p* = 0.52, one-way ANOVA; Fig. 5E). These results suggest that chronic alcohol exposure increases VTA-CeA connectivity, that this effect is measurable during both alcohol intoxication and acute withdrawal, and that this effect is not related to changes in intrinsic excitability of CeA neurons.

**Figure 5:**
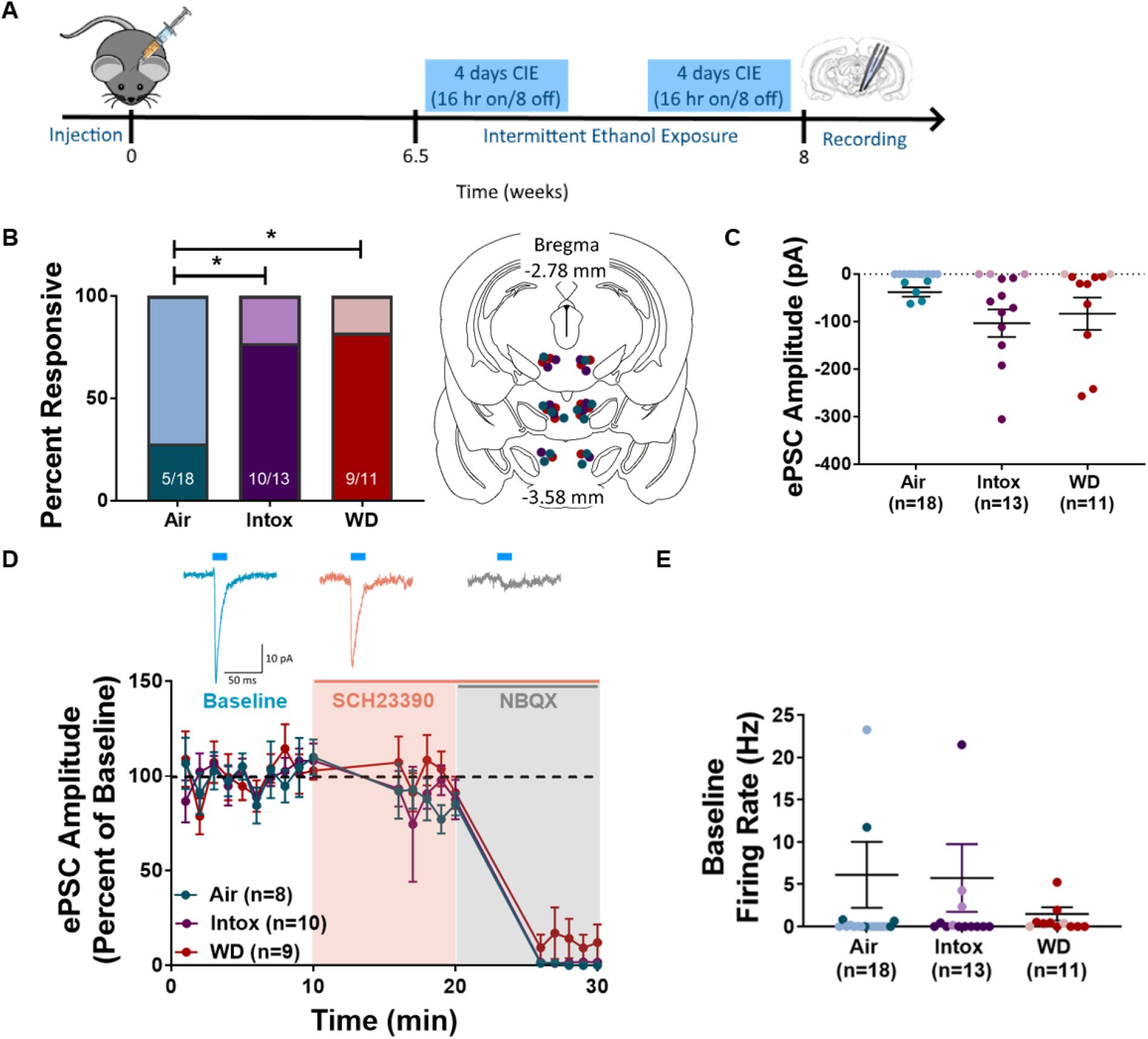
Proportion of CeA neurons responsive to stimulation of VTA terminals is significantly greater in alcohol dependent mice compared to naïve controls. **A**, Timeline of CIE experiments. **B**, Proportion of neurons in which an optically evoked event was elicited is significantly greater in dependent mice (either sacrificed during WD, or during intoxication) compared to naïve controls (\**p* < 0.05, Fisher’s exact test.) Inset shows center of VTA injection sites for dependent mice sacrificed during withdrawal (red), dependent mice sacrificed during intoxication (purple), or air controls (blue). **C**, Amplitude of ePSCs does not differ among groups. **D**, Time course of ePSC amplitude (as a percentage of baseline) in the presence of SCH 23390 (orange shaded area) and NBQX (grey shaded area). Representative baseline optically-evoked postsynaptic current (ePSC, blue) from stimulation of ChR2-containing VTA terminals in CeA (inset). Orange trace shows ePSC in the presence of SCH 23390; ePSC in the presence of SCH 23390 and NBQX shown in grey. Amplitude of ePSC (as a percentage of baseline) plotted over time. Data for Air group combines cells from air-exposed control mice with naïve mice from Fig. 3 (no difference in drug response was observed between groups). **E**, No significant difference in baseline firing rate of spontaneously active cells was observed when comparing cells across groups. Neurons in which an optically evoked-PSC was observed are indicated with darker data points. Line indicating mean firing rate was determined using spontaneously active neurons only. Data in **C**, **D**, and **E** shown as mean ± SEM.

## Discussion

The neural adaptations resulting from chronic alcohol use are not fully characterized, but may include a gradual recruitment of brain stress circuitry by mesolimbic reward circuitry that is activated during early stages of alcohol use. Here, we provide an anatomical and molecular characterization of a mesoamygdalar VTA-CeA circuit in two species commonly used in basic neuroscience. We demonstrate and map out a population of CeA-projecting VTA neurons in the rat, approximately one-third of which are dopaminergic, one-third glutamatergic, and one-third GABAergic. We also demonstrate the presence of VTA terminals in the CeA of mice and rats, noting spatial differences in the pattern of these projections between the two species. Optogenetic stimulation of VTA terminals elicits glutamate release in the CeA, a phenomenon which is more readily observed in the mouse than in the rat. Finally, we demonstrate withdrawal-induced activation of CeA-projecting VTA neurons in alcohol-dependent rats, and also chronic alcohol-induced increases in CeA neurotransmission from VTA terminals in alcohol-dependent mice. Collectively, these experiments increase our understanding of VTA-CeA circuit anatomy and physiology in rats and mice, and also demonstrate that alcohol withdrawal activates VTA-CeA cells in alcohol-dependent rats and mice, suggesting that this circuit may regulate alcohol dependence-associated behaviors.

Previous studies have characterized VTA projections to the CeA, but not extensively. Much focus has been on VTA DA neurons, as well as local modulation of DA signaling in the CeA. A population of CeA-projecting leptin receptor-containing VTA DA neurons was shown to regulate cocaine and amphetamine regulated transcript (CART) neurons in the CeA of mice^15^. In a separate study, optogenetic stimulation of DAT-expressing VTA terminals elicited glutamate release in 40% of CeA neurons recorded in *DAT::Cre* mice, indicating potential co-release of glutamate from VTA DA terminals in the CeA^14^. Here, we report a similar proportion of CeA neurons (33%) responding to optical stimulation of VTA terminals with an AMPA-sensitive postsynaptic current (Fig. 3). Rats and mice did not significantly differ in the proportion of optically-evoked events observed in the CeA after VTA terminal stimulation.

Characterization of the VTA-CeA circuit from anterograde or retrograde tracing, as outlined above, or from pharmacological manipulation within the CeA, has largely focused on the midbrain DA projections to CeA. For example, using genetic and pharmacological techniques in mice, one study showed that midbrain DA neuronal activity drives anxiety-like behavior that is reversed by intra-CeA administration of D1 antagonist SCH 23390^30^. Similar pharmacological approaches have been used to implicate CeA DA signaling in cocaine self-administration in the rat^31^. Here, we demonstrate that the CeA receives DA inputs from both the VTA and SN, suggesting that dopamine receptor pharmacology alone is insufficient to dissect the contributions of specific midbrain regions to dopamine-mediated effects in CeA (Fig. 1). We also demonstrate a substantial population of non-dopaminergic (glutamatergic and GABAergic) CeA-projecting midbrain neurons, suggesting that glutamate and GABA release from VTA terminals in the CeA may have important functional effects (e.g., on behavior). Indeed, a recent publication demonstrated a functional role for VTA-CeA GABAergic neurons in modulating defensive responses to a visual threat^16^. We observed functional VTA terminals in both the medial and lateral subdivisions of the CeA in the rat; it remains to be determined whether the functional or behavioral consequence of activation of these projections differs based on CeA subdivision. Activation of specific neuronal subtypes in medial and lateral CeA have differential behavioral responses (e.g., freezing, feeding), based on the projection target of these subpopulations^36–41^ (for review, see^42^). The CeA is not identical in mice versus rats, even at the anatomical level^35,50^, and there may be different connectivity patterns and function associated with different cell subsets in the two species. Future work is needed to fully examine VTA projections to CeA microcircuitry in rats and mice, which may be overlapping but not identical.

To our knowledge, no characterization of alcohol effects on VTA-CeA circuitry has been performed. There is some indirect evidence for a potential role of VTA-CeA circuitry in modulating alcohol drinking behavior. Orexin 1 receptor^43^ and nociceptin/orphanin FQ peptide receptor^44^ signaling in the VTA and CeA, as well as CRF binding protein and CRF 2 receptor signaling in VTA^45^, have been implicated in voluntary binge-like ethanol intake in mice and rats, although a direct connection between these regions has not been investigated in this context. Whether alcohol-induced increases in VTA-CeA activity is unique among drugs of misuse also remains to be determined. Dopamine in the CeA, presumed to come from the VTA, has been implicated in cocaine self-administration and withdrawal^31,46^, but not in nicotine withdrawal^47^. Chronic alcohol exposure increases neurotransmission in the CeA of alcohol-dependent rats during withdrawal^11^, which is likely to contribute to increased anxiety-like behavior and alcohol drinking in alcohol-dependent rats^13^. Unlike the CeA, which is often studied in the context of alcohol dependence, the VTA is more often associated with the initial, positive rewarding effects of ethanol^17^. While sub-chronic alcohol exposure increases the activity of VTA neurons (e.g.,^19,24^), chronic alcohol exposure is associated with decreased VTA activity or neurotransmission (e.g.,^48^). It is unknown whether early VTA engagement triggers plasticity in the CeA; here, we report alcohol withdrawal-induced activation of CeA-projecting VTA neurons in alcohol-dependent rats (Fig. 4). Only approximately one-third of these activated neurons express TH, suggesting that CeA-projecting glutamatergic and/or GABAergic VTA neurons also display increased activity during periods of withdrawal. Future work should better assess the identity of these neurons, although our data (Fig. 5) provide evidence for potential increases in excitatory input into the CeA from VTA terminals of alcohol-dependent mice, whether sacrificed during intoxication or during acute withdrawal.

Together, these data show that chronic alcohol exposure (i.e., alcohol dependence) leads to activation of the VTA-CeA circuit. Symptoms of AUD and alcohol dependence in humans and animals, such as heightened anxiety and alcohol intake, are often linked to CeA activity^49^, and given that data from transgenic mice provides indirect support for a role of VTA-CeA circuitry in mediating anxiety-like behavior^30^, we propose that activation of the VTA-CeA circuit may contribute to withdrawal-associated behaviors (e.g., increases in anxiety-like behavior and escalated alcohol drinking). Future work will test the role of the VTA-CeA circuit in mediating alcohol dependence-related behaviors.

## Acknowledgements

Supported by National Institute of Health grants R01 AA023305 (NWG), R01 AA026531 (NWG), R00 AA022651 (TAW), T32 AA007577 (EMA, CRK), F32 AA025831 (EMA). This work was also supported in part by Merit Review Award #I01 BX003451 (NWG) from the United States (U.S.) Department of Veterans Affairs, Biomedical Laboratory Research and Development Service.

## Author contributions

EMA & NWG designed the experiments; EMA, CRK, LKK, TDL, MC, TJT, TAW, & JWM performed research; EMA quantified data; EMA & NWG analyzed data; EMA & NWG wrote the paper; all authors edited the paper.

